# Determining the serotype composition of mixed samples of pneumococcus using whole genome sequencing

**DOI:** 10.1101/741603

**Authors:** James R. Knight, Eileen M. Dunne, E. Kim Mulholland, Sudipta Saha, Catherine Satzke, Adrienn Tothpal, Daniel M. Weinberger

## Abstract

Serotyping of *Streptococcus pneumoniae* is a critical tool in the surveillance of the pathogen and development and evaluation of vaccines. Whole-genome DNA sequencing and analysis is becoming increasingly common and is an effective method for pneumococcal serotype identification of pure isolates. However, because of the complexities of the pneumococcal capsular loci, current analysis software requires samples to be pure (or nearly pure) and only contain a single pneumococcal serotype. We introduce a new software tool called SeroCall, which can identify and quantitate the serotypes present in samples, even when several serotypes are present. The sample preparation, library preparation and sequencing follow standard laboratory protocols. The software runs as fast or faster than existing identification tools on typical computing servers and is freely available under an open source license at https://github.com/knightjimr/serocall. Using samples with known concentrations of different serotypes as well as blinded samples, we were able to accurately quantify the abundance of different serotypes of pneumococcus in mixed cultures, with 100% accuracy for detecting the major serotype and up to 86% accuracy for detecting minor serotypes. We were also able to track changes in serotype frequency over time in an experimental setting. This approach could be applied in both epidemiologic field studies of pneumococcal colonization as well as in experimental lab studies and could provide a cheaper and more efficient method for serotyping than alternative approaches.

## INTRODUCTION

*Streptococcus pneumoniae* (the pneumococcus) is a bacterial pathogen that causes a large burden of disease globally. Currently-available protein-polysaccharide conjugate vaccines target 13 of the more than 95 identified serotypes. The vaccines reduce the frequency of colonization due to vaccine-targeted serotypes and subsequently reduce disease [1]. There is a need to perform surveillance to monitor declines in vaccine-targeted serotypes as well as to detect increases in disease caused by serotypes not targeted by the vaccine (serotype replacement). The gold-standard is to monitor the incidence of invasive pneumococcal disease, a rare but severe outcome where the bacteria are isolated from a normally sterile site, such as the blood or cerebrospinal fluid. However, conducting such disease surveillance in low-resource settings is often not feasible. Therefore, it is often necessary to use other indirect measures of serotype epidemiology and vaccine effects. One such indirect measure is to track the prevalence of serotypes among healthy children who carry pneumococcus in the nasopharynx [2]. Because pneumococci are commonly detected among healthy children, point prevalence studies can be used to track changes in exposure to the different serotypes [3,4].

Carriage-based surveillance typically involves collecting a nasopharyngeal swab from a child, culturing it in the laboratory, isolating a pneumococcal colony, and then performing a traditional serotyping method such as the Quellung reaction, an antibody-based assay to determine the serotype of the isolate [5]. Quellung is relatively time consuming to perform, particularly when trying to test multiple colonies per sample. More recently, DNA based approaches have been used to determine the serotype of the isolated strain such as conventional and real-time PCR assays have been developed to identify common serotype and/or serogroups [6,7,8].

Whole-genome sequencing can effectively determine the serotype of single isolates, and several pipelines (PneumoCaT and SeroBA) have been developed [9,10]. A microarray-based platform can detect and quantify the relative abundance of all serotypes in a sample [11]. This is a highly-sensitive and accurate method and outperforms many other serotyping approaches [12]. The downside of this technology is that it requires specialized equipment that cannot be readily implemented by different labs. An ideal solution could be a sequencing-based approach that could be used to identify multiple serotypes in mixed samples and to quantify their abundance. Low-cost Illumina sequencing library preparation protocols make such an approach feasible and cost effective [13], and whole-genome sequencing is increasingly being adopted for diagnostic and public health applications [14]. The major challenge is a bioinformatic one: how to accurately identify and quantify serotypes in mixed samples.

The bioinformatic challenge revolves around the similarity of portions of the sequences in the capsular biosynthesis cassette in multiple serotypes. Only 25 of the 94 serotype capsular sequences are genetically distinct, while the rest form “serogroups” of genetically similar but phenotypically different serotypes [9]. The similarity is such that over 70% of error-free reads from those groups cannot uniquely map to a specific serotype, and several phenotypically distinct serotypes differ by only a single base pair over their 10-25 kb capsular sequence. Thus, traditional read mapping approaches fail, as they assume that nearly all informative reads will map uniquely. PneumoCaT and SeroBA can accurately identify serotypes from Illumina whole-genome sequencing reads. However, they expect “pure” samples (95% or more of the sample consists of a single serotype), do not provide quantitation, and will simply report “mixed” if the sample is found to contain multiple serotypes. In this study we develop and test an analysis approach and a software tool, SeroCall, for quantifying serotype abundance based on raw Illumina sequencing reads. We first use existing datasets and spiked samples in the lab to develop the pipeline. We then test the performance of this approach using a reference set of blinded gold-standard laboratory-prepared samples that have known quantities of different serotypes.

## METHODS

### Experiments with known serotype composition

The first set of dilution and competition experiments used invasive pneumococcal disease isolates that were obtained from the CDC’s Active Bacterial Core surveillance system isolate bank (1, 3, 4, 6B, 7F, 9V, 14, 18C, 19F, 23F). The strains were grown overnight on TSAII plates with 5% sheep’s blood at 37°C with 5% CO_2_.

#### Mixture with known concentrations of DNA

Overnight growth was harvested in PBS, and genomic DNA was extracted using a DNEasy blood and tissue kit (Qiagen) with the Gram positive pretreatment protocol. DNA was quantified using a NanoDrop reader (ThermoFisher). Serotypes were then mixed together at the following ratios: 19F:23F[1:2], 19F:23F[1:4], 19F:23F[1:8], 19F:23F[1:10], 19F:23F[1:100], 19F:23F:1 [1:4:1],19F:23F:1:4:18C [1:2:4:2:1]. These samples were sequenced to an average of 2.76 million reads per sample on an Illumina HiSeq.

#### Longitudinal growth experiment

Overnight growth on TSAII plates was resuspended in PBS, and the optical density (OD) at 600nm was adjusted to 0.05. These stocks were then diluted 1:20 into a diluted broth of 7.5mL PBS, 2.5mL BHI, 8.25uL sheep blood, and 125uL horse serum. The strains were grown individually for several hours until the lowest concentration strain reached OD ~0.15. All strains were adjusted down to match this value. The strains were then mixed at equal concentration and diluted 1:20 into fresh broth in deep well plates (800 uL broth + 40uL bacteria) with a separate replicate well for each time point. The plate was then incubated at 37°C with 5% CO_2_. 40uL from each well of the 6h time-point was used to seed a new row of 800uL broth to allow another 2 hours of growth. This passaging step is important because in the limited nutrient broth, pneumococcal population tends to crash after 6 hours. At the indicated time points, the full volume of the well was transferred to the −80 °C freezer. At the end of the experiment, DNA from all wells was extracted at the same time using a Qiagen DNEasy blood and tissue kit with Gram positive pretreatment protocol (Qiagen). The experiment was performed in duplicate.

### Single serotype calls and benchmarking against other serotyping software

The development and validation whole-genome sequencing datasets that were used in [9] to develop and test the PneumoCaT and SeroBA software were used similarly here. 871 development samples, covering all 94 serotypes, were used in the development of the SeroCall software, and then the 2065 validation samples covering 72 serotypes were evaluated using the final version of the software. Also, the current versions of PneumoCaT (v1.2) and SeroBA (v1.0.1) were run locally on all samples, and the “gold standard” calls used for benchmarking were the majority vote of the original laboratory serotyping, the PneumoCaT call and the SeroBA call. This accounts for updates to the software that correct for serotypes reported at the time of the PneumoCaT publications (most notably, a change in the 12B vs. 12F typing that was discovered after the publication of [9]).

### Evaluation with blinded samples

The PneuCarriage Project [12] was a multi-center study to evaluate pneumococcal serotyping methods. A set of standard laboratory-prepared sample mixtures has been evaluated using a large number of serotyping methods. Eighty of these samples, containing mixtures of 0-4 serotypes, were evaluated using our analysis pipeline. The laboratory personnel processing the samples and the bioinformatic analysts were blinded to the serotype composition of the samples.

The samples were provided as frozen aliquots. Samples were thawed, 10-fold dilutions were spread on a TSAII plate with 5% sheep’s blood, and incubated overnight at 37°C with 5% CO_2_. The most concentrated non-confluent dilution was harvested into PBS, and DNA was extracted as described above. During the culturing and preparation, 9 of the serotype-positive samples failed to culture, and a further 6 samples failed amplification due to an existing primer failure. The remaining 65 samples were sequenced to an average of 1.90 million reads per sample; a second round of sequencing increased the average reads per sample to 4.67 million reads per sample. Serotype calls and quantifications were returned for all samples and evaluated by PneuCarriage Project personnel.

### Library preparation

Illumina libraries were prepared using the protocol described by Baym, *et. al.* [13]. The exception was that the final cleanup was performed using Qiagen PCR purification columns rather than the bead-based assay described in the original paper. The sequencing indices that were used are listed in the supplemental material. Each plate of up to 96 samples were multiplexed and run on an Illumina HiSeq with a read length of 2×150bp.

### Bioinformatic approach

The overall steps of the SeroCall software follow that of PneumoCaT and SeroBA, but apply different algorithms in order to quantify all serotypes found in a sample. All of the tools (1) align the read data to the set of serotype capsular sequences, (2) identify serogroups and distinct serotypes that are present in the sample, and (3) distinguish serotypes within the serogroups, using serotype-specific variants or regions of the capsular sequences.

#### Step 1 Read Alignments

Sequence read data are first aligned using BWA MEM [15] to a “reference” that combines the serotype capsular sequences from 94 serotypes with the non-capsular sequences from three *S. pneumoniae* genomes: R6, SPNA45 and ATCC700669 (where the capsular sequence from each has been masked). The supplementary material details the capsular reference sequences used, most of which are the same as the PneumoCaT reference sequences, but several have been modified to work with this algorithm. The genome sequences mainly serve as a “decoy”, so that genomic reads will align to the genome sequences instead of to the capsular sequences, and so will not affect the read depths for the serotypes.

The read alignments produced by BWA MEM are used to compute “bin counts,” counting the total and uniquely-mapped reads across the serotype capsular sequences. Each sequence is partitioned into 500 bp bins, denoted S_i,b_ for serotype i and bin b. The choice of 500 bp ensures that local depth variations resulting from sequencing are smoothed in the bin counts. Read alignments are processed in read-pairs, and are first filtered for (1) any genomic alignments, (2) any unmapped alignments (if either read in a read pair is unaligned, then both reads in the pair are filtered), (3) any chimeric reads aligning to two different serotypes, or (4) any read pairs with a combined 10 or more differing or soft clipped bases, as this is a sign of a genomic read pair mistakenly aligned to a serotype sequence.

The remaining read-pair alignments add to bin counts, incrementing the counts of any bin which overlaps with either of the read alignments. The “total bin counts” count both uniquely mapping reads (reads whose MQ >= 0) and repetitively mapping reads (could map equally well to multiple locations in the serotype sequences, where the BWA MEM software randomly chooses a location from those best locations). The “unique bin counts” count only uniquely mapping reads. For an input sequence dataset, this results in T_i,b_ and U_i,b_ matrices containing the bin counts for that data.

#### Step 2 Serotype/Serogroup Quantification

The second step takes the bin counts from the input data, treating them as the “observed” counts OT_i,b_ and OU_i,b_, and compares them against the sets of “expected” counts ET_y,i,b_ and EU_y,i,b_ for all serotypes y (because of the similarity between serotype sequences, reads from a serotype y will align to serotype i, and so will contribute to the bin counts for serotype i). These expected counts were determined by generating simulated reads for each serotype and computing the bin counts for those reads. Specifically, a simulated 2×100bp read-pair (with an insert size of 200 bp) was generated at every position of a serotype’s capsular sequence, so that each location of the capsular sequence is covered by 200 reads across the whole sequence, except at the ends. Generating bin counts in this way results in expected bin counts for an equal 200x sampling of each serotype.

The comparison uses expectation-maximization to compute the optimal “factor levels” F_y_ for each serotype y, which optimize the equations:

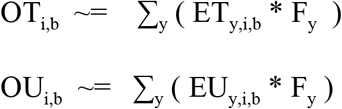

In other words, it computes the factor levels that result in a mixture of the expected serotype bin counts which most closely resembles the observed bin counts. Initially, all F_y_ are set to 1.0, then 100 rounds of a gradient descent algorithm is performed, computing the expected mixture bin counts, comparing them to the observed counts and adjusting the factor levels up or down.

Specifically, each round of the algorithm first computes MT_i,b_ = ∑_y_ (ET_y,i,b_ * F_y_) and MU_i,b_ = ∑_y_ (EU_y,i,b_ * F_y_), each across all serotypes y, then it computes bin ratios RT_i,b_ = OT_i,b_ / MT_i,b_ and RU_i,b_ = OU_i,b_ / MU_i,b_. Ideally, all of the ratios should equal 1.0, if the computed mixture matches the observed counts across all of the bins. Per-serotype factor ratios, ratioT_i_ and ratioU_i_, are then computed using a weighted median of the bin ratios for the serotype, where the weights for each bin are ET_i,i,b_ / maxET and EU_i,i,b_ / maxEU (i.e., the fraction of total/unique reads coming from serotype i and bin b that were actually counted in the bin). This gives higher weight to the more unique, or less repetitive, regions of each serotypes’ capsular sequence. And, the median is used instead of the mean in order to prevent genome contamination and local genetic differences (serotype samples whose actual capsular sequence mainly matches the reference, but which contains a local region unique to another serotype) from skewing the quantitation.

Final serotype ratios are computed by combining the ratioT and ratioU values based on the bin with the highest unique weight, i.e., mW_i_ = max_b_ (EU_i,i,b_ / maxEU). And so, ratio_i_ = mW_i_ * ratioU_i_ + (1.0 - mW_i_) * ratioTi. Then, new factor levels are computed as F_i_ = F_i_ + (F_i_ - F_i_ * ratio_i_) / 2, adjusting the factor level by half of the computed observed over expected ratio, in each round of the gradient descent. Also, if at any point, F_i_ falls below 0.002, it is set to 0.0.

#### Step 3 Serogroup Refinement

The third and final step readjusts the factor levels for the serotypes within serogroups that cannot be distinguished based on bin-sized read depth differences. The method of Steps 1 and 2 are able to distinguish 56 of the serotypes without further refinement, and the groups of serotypes that require further refinement by this method are: 6A/6B/6C/6D/6E; 7A/7F; 7B/40; 9A/9V; 11A/11D; 11B/11C; 12A/12B/12F/44/46; 15B/15C; 18B/18C/18F; 24B/24F; 25A/25F/38; 32A/32F; 33A/33F/37; 35A/35C/42.

Serotypes 15B and 15C can interconvert [16] and so are reported as 15B/15C. In keeping with the reporting performed by PneumoCaT, the serogroups 24 and 32 are reported at the group level. For the other groups, the CTV database in PneumoCaT was used to identify variants and genes/alleles that differ between serotypes in the groups. Each of those differences was translated into the sets of locations within the full capsular sequence references, instead of the CTV database’s reporting of a gene position or gene sequence. For example, the CTV database identifies a SNP in *wcjE* at position 721 where serotypes 9A and 9V are different. This is translated into the locations 18,534 and 18,852 in the 9A and 9V capsular sequences, respectively. The list of all differences used is given in the supplementary material.

The reason for the translation is that this step of the algorithm takes advantage of the sensitivity of BWA MEM in aligning reads to near-identical locations in the reference. If there is a single location whose alignment contains more identities than all other locations, that “best match” location will be chosen. Even a single nucleotide difference is sufficient for BWA MEM to consistently align reads to the proper serotype’s sequence, and so the read depths at those difference locations provide an accurate measure of the differences between serotypes. So, instead of performing a separate variant calling, mapping or *de novo* assembly to resolve serotypes, the computation of Step 1 computes “bin” counts at these specific difference locations, and then this step compares the observed counts at those locations against the expected counts.

Since these locations are where the serotypes are genetically different, and alignment “bleed” is not an issue, this step just computes depth ratios for each serotype and difference location in a group, where OU / EU is used if EU is greater than 0, and OT / ET is used otherwise. The ratio for a serotype is the minimum of the computed depth ratios across the difference locations (if the serotype is present in the sample, each of these locations should have a non-zero read depth). Then, those ratios are summed, and serotype percentages are computed by dividing the serotype ratio by the sum of the ratios.

If the sum of the ratios is 0, this means that there is no read evidence distinguishing the serotypes in the group, and an ambiguous call like “09A/09V” is made, with a factor level equal to the sum of the factor levels computed in step 2, for the serotypes in the group. If the sum is greater than 0, then the factor levels from step 2 (again, for the serotypes in the group) are reapportioned using the serotype percentages computed in this step. So, for example, if 09A and 09V had step 2 factor levels of 0.13 and 0.11, but the step 3 serotype percentages were 80% 09A and 20% 09V, then the factor levels would be changed to 0.192 for 09A and 0.048 for 09V (to maintain the 09A/09V levels compared to all other serotypes, but reset the serotype-specific levels to the identified percentages).

Once the final factor levels are computed, they are converted to percentages by dividing each by the sum of all factor levels. Then, any serotype with a percentage less than 0.2% is filtered out, and the percentages are recomputed using only the remaining serotypes. Those serotypes and percentages form the output calls produced by the software.

#### Data and software availability

The PneumoCaT serotype datasets can be accessed through the European Nucleotide Archive (ENA) under project PRJEB14267. The known mixture and replicate sample datasets can be accessed through the NCBI Sequence Read Archive under project PRJNA561126. The software is freely available under an open source license at https://github.com/knightjimr/serocall.

## RESULTS

### Single serotype calls using sequences in the PneumoCaT database

For the 871 development samples, the calls made by SeroCall had a concordance rate of 96.1% at the serotype level and 98.6% at the serogroup level, with 816 exactly matching calls, 21 samples with only minor differences (i.e., a matching serotype call with abundance above 95%, which is the PneumoCaT threshold for reporting a “pure” serotype, plus additional calls with total abundance below 5%), 22 samples with multiple or different serotypes called from the matching serogroup, and 12 “discrepant” samples.

For the 2065 validation samples, the concordance rate was 97.0% at the serotype level and 98.9% at the serogroup level, with 1924 exactly matching calls, 79 samples with minor differences, 40 samples with multiple/different serotypes from the matching serogroup called, and 22 “discrepant” samples. The details of all non-matching samples for both datasets can be found in the supplementary material.

The average, minimum and maximum execution times for SeroCall, SeroBA and PneumoCaT are given in Table 1, for the analysis of the development samples (the validation sample running times were similar). All software was run on 20 core, 121 GB memory, “2x E5-2660 v3” compute servers, where the software was run with exclusive access to the server. The SeroCall and PneumoCaT command lines were passed “-t 20” options, allowing them to use 20 parallel threads. For this dataset on these servers, SeroCall ran three times faster than SeroBA and twice as fast as PneumoCaT. The computation in SeroCall is dominated by the BWA MEM alignment, which scales linearly in the number of cores. So, on compute servers with 8 or more cores, SeroCall is expected to run as fast or faster than SeroBA.

**Table 1.** Comparison of run times for SeroCall, SeroBA, and PneumoCAT.

| Running time (MM:SS) | SeroCall | SeroBA | PneumoCaT |
| --- | --- | --- | --- |
| Minimum | 0:14 | 0:26 | 0:17 |
| Average | 0:35 | 1:43 | 1:16 |
| Maximum | 1:06 | 2:39 | 2:49 |

### Mixed samples with known concentrations

Mixtures of DNA were prepared, containing known fractions of 2, 3 and 5 different serotypes. SeroCall was able to accurately recover the true fraction of each serotype, including serotypes that were present at a low fraction (Figure 1).

**Figure 1.**
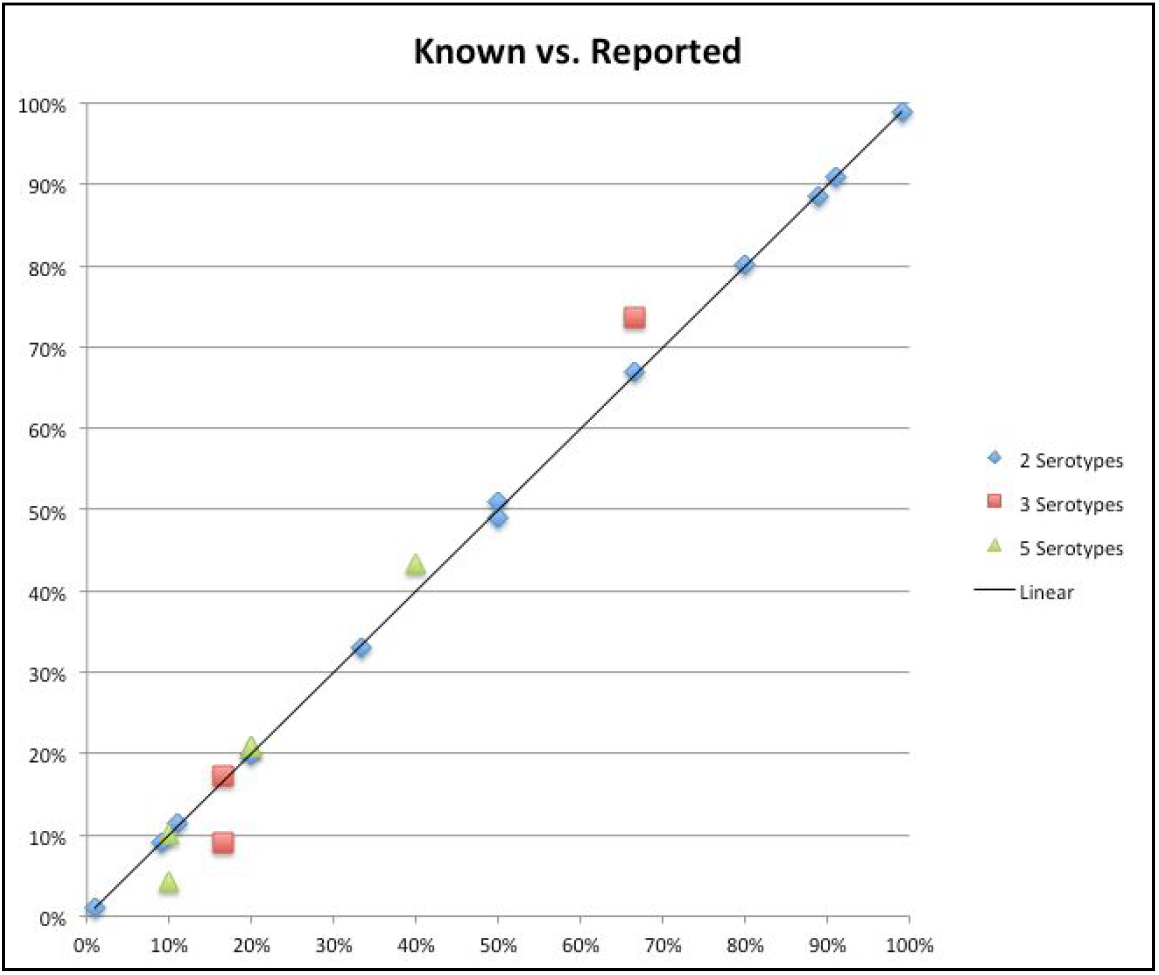
Comparison of the true and estimated percentage of each serotype in samples with 2, 3, and 5 serotypes.

#### Blind testing of mixed serotype samples from the PneuCarriage project

Fifteen of the 80 PneuCarriage samples either failed to grow (n=9) or had other technical issues during library preparation (n=6). In keeping with the blind testing, results from all samples were returned to the PneuCarriage project and evaluated. Here we outline the results of the 65 samples that were successfully cultured, prepared and sequenced. The supplementary material details the full evaluation of all 80 samples.

Table 2a shows the sensitivity of the assay, both with a first round of sequencing, along with a second round of sequencing that increased the average reads per sample. The sensitivity to detect the major serotype was 98% and 100% for the first and second rounds of sequencing, respectively. Samples containing serotype 12F were misidentified as 12B, as the version of SeroCall used in this testing was based on the original CTV database from the PneumoCaT paper [9]. The sensitivity to detect minor serotypes was 59% using only the 1.9 million reads per sample (first round), and improved to 81% with 4.67 million reads per sample (second round). However, that came at the cost of an increase in false positive identifications. Excluding 12F/12B misidentifications, there was one false positive in the first round (resulting in a positive predictive value (PPV) of 96%) but six false positives in the second round (PPV of 95%).

**Table 2.**
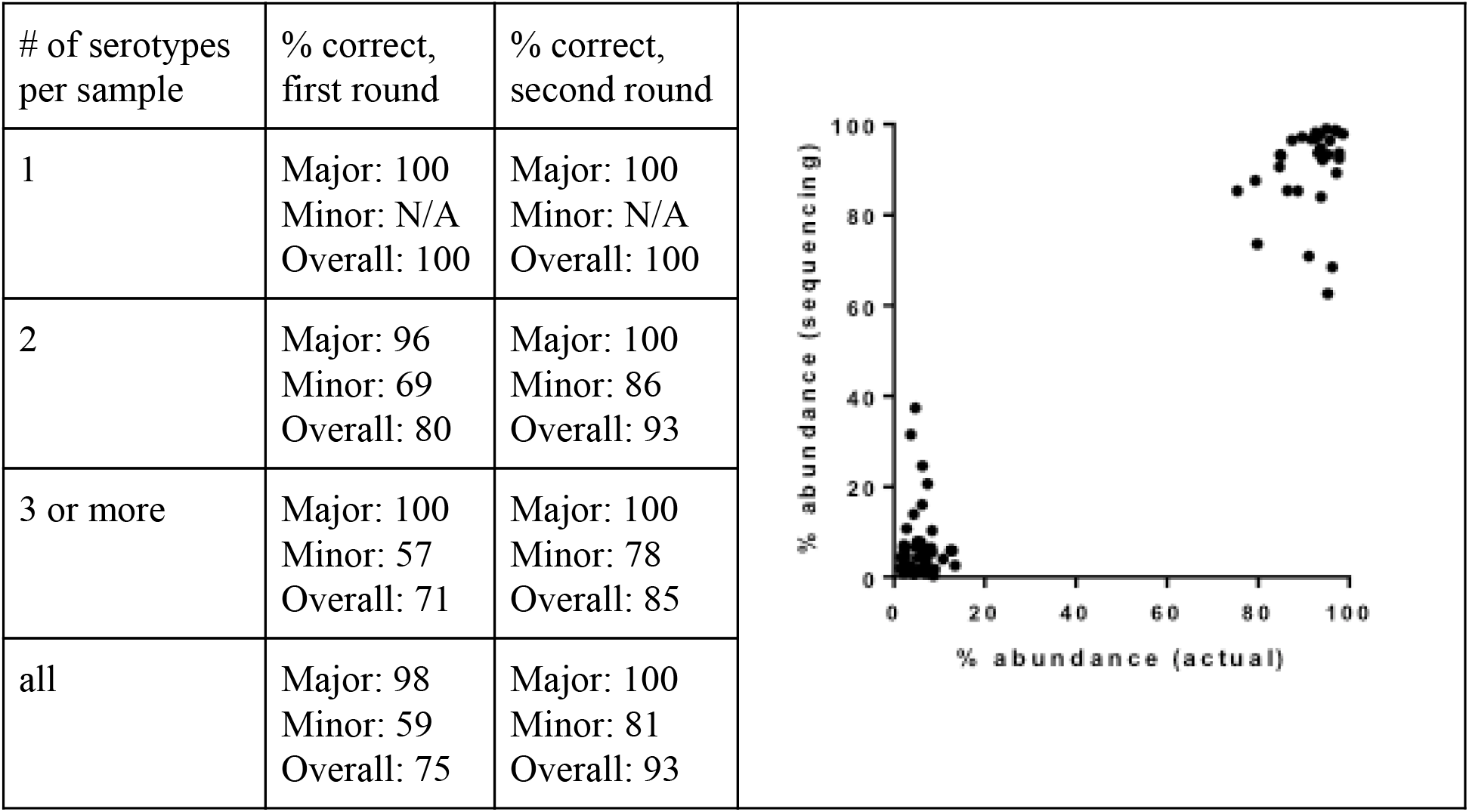
(a) Serotype calling accuracy for the 65 PneuCamiage blind testing samples, using first round of 1.9 million reads per sample and second round using 4.6 million reads per sample. (b) Comparison of SeroCall quantification ("abundance (sequencing)") and known PneuCaniage abundances ("abundance (actual)"), for 32 mixed samples.

Finally, the quantitation of the serotype calls was evaluated against the known spiked levels for 32 multi-serotype samples called correctly (using the second round data). The correlation between the two was strong (Spearman’s p = 0.762, p < 0.0001), and the mean absolute difference between the known level and the SeroCall quantitation was 4.0% (Table 2b).

#### Longitudinal monitoring of mixed samples and replicates

Finally, we sought to evaluate the ability of this approach to track changes in serotype frequency over time, approximating the setup with longitudinal carriage sampling. Mixtures of 2 to 10 clinical isolates, representing different serotypes, were grown *in vitro* (in duplicate) and sampled at 2, 4, 6 and 8 hours. These replicate samples were prepared and sequenced, testing both longitudinal monitoring of changes in serotype frequency and testing reproducibility. There was good agreement between replicate samples, and it was possible to track changes in the frequency of individual serotypes over time (Figure 2). Note, because of a primer failure, only single results were generated for the 10 serotype sample at 6 hours and 8 hours.

**Figure 2.**
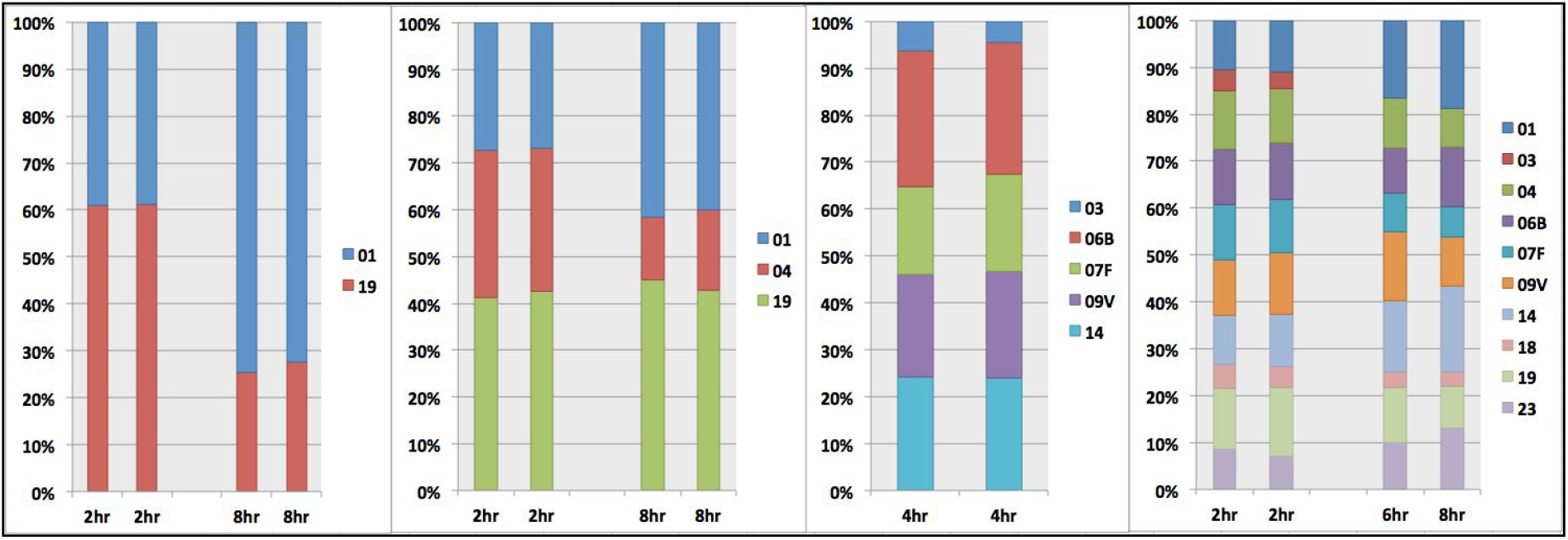
Replicate testing using mixtures of 2, 3, 5 and 10 serotypes. Replicates cultured for 2, 4, 6 or 0 hours before selection for sequencing.

## DISCUSSION

We developed and validated a whole-genome sequencing method and analysis software SeroCall for the identification and quantification of *Streptococcus pneumoniae*. The software was tested using both internally and externally generated samples and datasets, and returned concordant results with other serotyping methodologies and other WGS-based serotype identification software. It also performed well in tracking changes in the proportion of serotypes over time in mixed cultures in an experimental setting. SeroCall is the first sequencing-based method to perform serotype quantitation on mixed samples. Its computational performance matches or exceeds that of other software, and could be implemented efficiently to enable high-throughput surveillance of population serotypes.

Antibody-based serotyping (Quellung and latex-agglutination kits) are easy and fast when checking the serotype of one colony. However, understanding multiple carriage is difficult with Quellung [17]. Scaling to large number of samples can also be labour intensive with antibody-based methods. Existing DNA-based methods have insufficient sensitivity or specificity to identify multiple serotypes, or are microarray based. Compared to microarrays, access to/use of sequencing is increasing for various public health surveillance, and costs are going down. So implementing carriage surveillance with a low-cost sequence-based multiple carriage surveillance is timely.

While the method was concordant with other methodologies, there are still some limitations. As for all serotyping methods, upstream aspects including sample storage and culture remain important; this is reflected in the fact that we could not obtain results from nine culture-negative samples from the PneuCarriage set. The *in vitro* culture step prior to DNA extraction, as well as the DNA extraction protocol itself, could influence the detected proportion of serotypes. The quantification methods use the current set of capsular sequences as a core “truth set” from which to compare the sequencing results. If there are CPS biosynthesis loci recombinations that are not present in the current dataset, serotypes might be misquantified or misclassified. SeroCall uses the entire genetic sequence to perform is classifications and quantifications, and for samples where the phenotypic serotype differs than the overall genetic serotype ancestry, misclassifications may occur. Also, complex mixtures of very closely related serotypes may be more difficult to accurately quantify than more genetically distinct serotypes. Additionally, non-pneumococcal streptococci present in the nasopharynx can confound sequenced-based serotyping [18]. As such, future work will evaluate SeroCall using nasopharyngeal samples, for example the PneuCarriage field samples (aliquots of STGG from NP swabs). Finally, improvements and increased testing of the laboratory methodologies, including the use of non-duplicate Illumina index primers and potentially side-by-side comparison of microarray and sequencing of the same DNA, to explore differences in DNA extraction efficiency, may increase the robustness of the method.

One experimental parameter that affects the results is the read depth. Using the blinded samples, we found that with a low read depth, the sensitivity to detect minor serotypes was greatly reduced. We recommend obtained a read depth of 2-3 million reads per sample to obtain sensitivity similar to what is reported here, and possibly higher depth if looking for very low abundance serotypes. The ‘cost’ of increased read depth was an increase in the number of false-positive identifications of rare serotypes. One possible explanation for the false positives is Illumina “barcode hopping” [19], as the library preparation used standard multiplex primer sequences which have been found to be susceptible to that. Using unique barcode pairs for each sample could help to avoid this issue and allow for detection of low abundance serotypes without an increase in false positivity.

The library preparation protocol that we use, which was developed by Baym et al. [13], can produce high-quality, low-cost sequences when multiplexing samples. Provided that an investigator has access to an Illumina sequencer, this makes performing sequence-based serotyping cost effective when compared with traditional serotyping methods.

During sample preparation we lost several samples. 9 of the blinded samples failed to culture. This could have been an issue with sample transport or with the culture conditions in the lab. We also lost several samples due to a primer failure during the library preparation. Confirming the concentration of each sample prior to pooling would catch this issue prior to pooling to allow for re-generation of the libraries for the affected samples.

In conclusion, we describe the development of an analytical tool that can be used to quantify the abundance of multiple serotypes in mixed cultures using a sequencing-based approach. We do this by addressing a bioinformatic challenge in assigning Illumina reads from a mixed sample to the correct serotype. This method could be applied to epidemiologic studies of pneumococcal carriage that seek to evaluate the carriage frequency of dominant and sub-dominant serotypes and can be used to monitor changes associated with the introduction of conjugate vaccines.

## Supporting information

S1. Discrepant calls for PneumoCaT samples

## Acknowledgements

Invasive disease isolates were obtained from the Active Bacterial Core surveillance (ABCs)/Emerging Infections Programs (EIP) Network. Thank you to the PneuCarriage project group [12] for their provision of well-characterised spiked samples for testing in this project.

## SUPPLEMENTARY MATERIAL

### Reference sequences used by SeroCall

The capsular sequences from PneumoCaT’s v1.2 streptococcus-pneumoniae-ctvdb database were used as the reference capsular sequences for SeroCall, with the following alterations:

- PneumoCaT’s 06E sequence was duplicated into 06E_6A and 06E_6B, and then the 06E_6A sequence was edited, changing base 7912 from A to G (to match the difference used to distinguish 6A and 6B, see below).
- PneumoCaT’s 25A sequence was edited so that the *wcyC* sequence matches the *wcyC*_25A sequence found in 25A_25F_38/reference.fasta. The reason for this is that the PneumoCaT 25A sequence, and the original 25A reference from [1] (accession CR931689.2), contain the *wcyC*_25F sequence, not *wcyC*_25A. (This does not affect PneumoCaT or SeroBA, because they use the separate gene sequences in the second phase of their analysis. However, SeroCall uses only the primary reference, so it requires the correct 25A *wcyC* sequence to be there.)
- PneumoCaT’s 33F sequence was extended with the rest of the sequence from [1] (accession CR931702.1), starting at base 1205 of that sequence, so as to restore the presence of the *wcjE* gene in the capsular sequence. (The 14,496 bp PneumoCaT sequence exactly matches CR931702.1 from base 1205..15700, but does not contain the last 1,299 bases of CR931702.1.)
- PneumoCaT’s 37 sequence, which consists only of the tts gene sequence, was replaced with the sequence from [1] (accession CR931709.1) starting at base 1587, which is the location matching the beginning of the PneumoCaT 33A and 33F sequences. Then, that sequence was appended by 40 N’s and the *tts* gene sequence. This was done so that reads from the non-functional 37 capsular sequence are aligned to this sequence, instead of being aligned to the 33A and 33F sequences.

The *S. pneumoniae* genome sequences used as decoy sequence for read alignments are the R6 reference (accession NC_003098.1), SPNA45 reference (accession HE983624.1) and ATCC700669 reference (accession FM211187.1).

### Sequence differences used to distinguish serogroup members

In phase 3 of the SeroCall algorithm, sets of locations are used to distinguish the presence of specific serotypes that cannot be distinguished by larger genetic differences in phase 2. The following difference locations are used:

| ID | Serotype:Location |  |  |  |  |
| --- | --- | --- | --- | --- | --- |
| diff1 | 06A:7577 | 06B:7577 | 06C:7384 | 06D:7384 |  |
| diff2 | 06A:5400-5500 | 06B:5400-5500 | 06C:5400-5500 | 06D:5400-5500 |  |
| diff3 | 06E(6A):7850 | 06E(6B):7850 |  |  |  |
| diff4 | 07A:8598 | 07F:8582 |  |  |  |
| diff5 | 07B:8250 | 40:8259 |  |  |  |
| diff6 | 07B:8590 | 40:8600 |  |  |  |
| diff7 | 09A:16996 | 09V:16996 |  |  |  |
| diff8 | 11A:8237 | 11D:8237 |  |  |  |
| diff9 | 11B:13140 | 11C:13122 |  |  |  |
| diff10 | 12A:4900-5000 | 12B:4200-4300 | 12F:4200-4300 | 44:4200-4300 | 46:4900-5000 |
| diff11 | 12A:8077 | 12B:7195 | 12F:7204 | 44:7204 | 46:8077 |
| diff12 | 12A:9822 | 12B:8940 | 12F:8949 | 44:8949 | 46:9822 |
| diff13 | 12A:12004 | 12B:11122 | 12F:11131 | 44:11131 | 46:12004 |
| diff14 | 12A:12217 | 12B:11335 | 12F:11344 | 44:11344 | 46:12217 |
| diff15 | 18B:7700-7800 | 18C:7700-7800 | 18F:7700-7800 |  |  |
| diff16 | 18B:12382 | 18C:12382 | 18F:12382 |  |  |
| diff17 | 25A:11800-11900 | 25F:11800-11900 | 38:10000-101100 |  |  |
| diff18 | 25A:10477 | 25F:10477 | 38:8645 |  |  |
| diff19 | 28A:7049-7080 | 28F:7049-7080 |  |  |  |
| diff20 | 33A:14968 | 33F:14968 | 37:16400-16500 |  |  |
| diff21 | 35A:10753 | 35C:10758 | 42:10758 |  |  |
| diff22 | 35A:1973 | 35C:1982 | 42:1982 |  |  |

### Differences in calls made using the PneumoCaT Development and Validation databases

The supplemental Excel spreadsheet lists the development and validation samples where the SeroCall result differed from a 100% call of the majority call made by serotyping from [2], PneumoCaT and SeroBA.

### Full sensitivity table from the PneuCarriage report

Following the reporting standards used by the PneuCarriage project, all samples tested were used to determine the sensitivity and specificity of the serotyping method. This included samples that were successfully sequenced as well as those that failed to culture and those that had other sample preparation failures. The table below was taken directly from the PneuCarriage report, giving the full sensitivity metrics (1) for all samples, (2) for all samples except the 9 culture negative samples, and (3) for all samples except the culture negative and primer failure samples. Also, the table reports results from both rounds of evaluation, where the first round occurred after one round of sequencing (with an average of 1.9 million reads per sample), and the second occurred after a second round of sequencing was performed (increasing the average read count to 4.65 million).

|  | First Round Results |  |  | Second Round Results |  |  |
| --- | --- | --- | --- | --- | --- | --- |
| Number of serotypes per sample | % correct | % correct (excluding 9 culture failures) | % correct (excluding 15 culture, primer failures) | % correct | % correct (excluding 9 culture failures) | % correct (excluding 15 culture, primer failures) |
| 1 | Major: 71<br>Minor: N/A<br>Overall: 71 | Major: 83<br>Minor: N/A<br>Overall: 83 | Major: 100<br>Minor: N/A<br>Overall: 100 | Major: 71<br>Minor: N/A<br>Overall: 71 | Major: 83<br>Minor: N/A<br>Overall: 83 | Major: 100<br>Minor: N/A<br>Overall: 100 |
| 2 | Major: 73<br>Minor: 49<br>Overall: 61 | Major: 84<br>Minor: 56<br>Overall: 70 | Major: 96<br>Minor: 69<br>Overall: 80 | Major: 76<br>Minor: 65<br>Overall: 70 | Major: 88<br>Minor: 75<br>Overall: 81 | Major: 100<br>Minor: 86<br>Overall: 93 |
| 3 or more | Major: 88<br>Minor: 49<br>Overall: 61 | Major: 96<br>Minor: 55<br>Overall: 68 | Major: 100<br>Minor: 57<br>Overall: 71 | Major: 88<br>Minor: 67<br>Overall: 74 | Major: 96<br>Minor: 76<br>Overall: 82 | Major: 100<br>Minor: 78<br>Overall: 85 |
| all | Major: 79<br>Minor: 49<br>Overall: 61 | Major: 88<br>Minor: 55<br>Overall: 70 | Major: 98<br>Minor: 59<br>Overall: 75 | Major: 80<br>Minor: 66<br>Overall: 72 | Major: 90<br>Minor: 76<br>Overall: 82 | Major: 100<br>Minor: 81<br>Overall: 93 |

### Sequencing indices used for multiplexing

Samples from 96-well plates were barcoded using standard Illumina multiplexing PCR primers,

Index 1 Read: `5′ CAAGCAGAAGACGGCATACGAGAT[i7]GTCTCGTGGGCTCGG`

Index 2 Read: `5′ AATGATACGGCGACCACCGAGATCTACAC[i5]TCGTCGGCAGCGTC`

where the following i5 and i7 indices were used for the samples in each well:

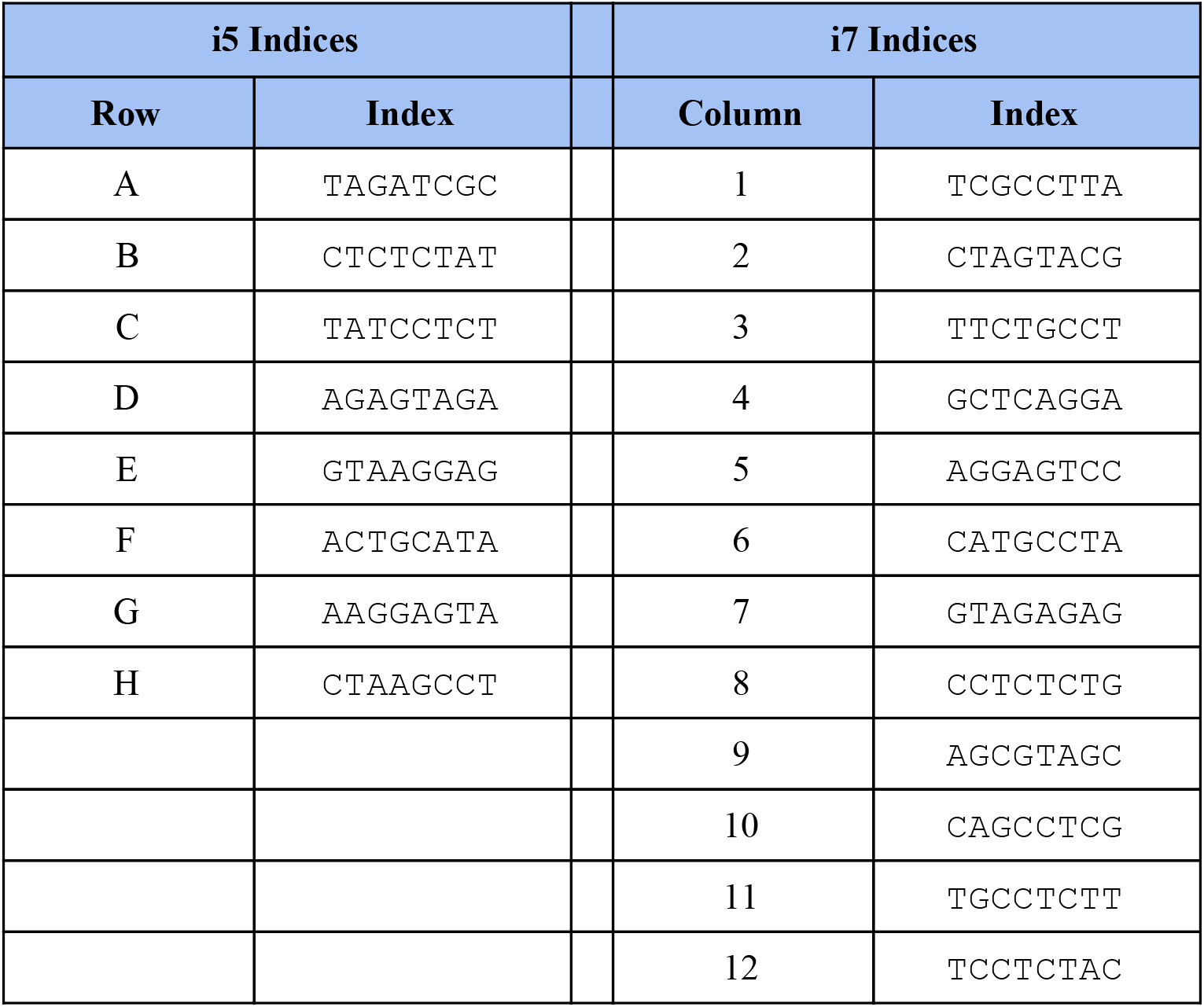

